# GIP: A Gene network-based integrative approach for Inferring disease-associated signaling Pathways

**DOI:** 10.1101/654780

**Authors:** Xi Chen

## Abstract

Dysregulation or crosstalk of signal transduction pathways contributes to disease development. Despite the initial success of identifying causal links between source and target proteins in simple or well-studied biological systems, it remains challenging to investigate alternative pathways specifically associated with a disease. We develop a <u>G</u>ene network-based integrative approach for <u>I</u>nferring disease-associated signaling <u>P</u>athways (GIP). Specifically, we identify alternative pathways given source and target proteins. GIP was applied to human breast cancer data. Experimental results showed that GIP identified biologically meaningful pathway modules associated with antiestrogen resistance.

## Introduction

To address the complexity and heterogeneity inherent in most cancers, effective combinatorial drug design requires the development of novel systems biology tools to identify cancer-specific signaling pathway. Given the importance of understanding the complex signal transduction pathways, computational and integrative methods continue to explore new means to effectively integrate multi-platform genomic data with biological knowledge. Gene Set Enrichment Analysis (GESA) ^1^ and PAthway Recognition Algorithm using Data <u>I</u>ntegration on Genomic Models (PARADIGM) ^2^ are useful tools being able to capture disease-specific activities of canonical pathways. Yet, their ability to discover novel pathway interactions remains debatable.

Growing knowledge about genome-wide protein-protein interactions (PPIs) ^3,4^ has offered an alternative source of information for signaling network identification ^5^. One major challenge in pathway analysis is to infer edge directions based on non-directed PPI interactions. Among limited methods ^6,7^ that specifically tackle the problem of inferring edge directions for pathway identification, most of them either reply on protein domain interactions or are not scalable to the entire proteome. Notably, Gitter *et al.* proposed to use maximum edge orientation (EO) on PPI network to determine the most likely signaling directions that fulfil global optimality ^8^. However, the EO approach relies heavily on the assumption that most biological pathways are short (length<5) in order to accommodate the requirement of exhaustive enumeration of all possible pathways. Neither does EO take into account important biological knowledge such as subcellular information, which often yields signal transduction directions that are difficult for biological interpretation.

To reconstruct aberrant pathway modules in cancer studies, we develop a <u>G</u>ene network-based integrative approach for <u>I</u>nferring disease-associated signaling <u>P</u>athways (GIP). GIP integrates gene expression data with PPI networks in a distribution learning framework to study aberrant signal transduction and alternative pathways. The core functionalities of GIP are built upon two functional modules: (1) a Gibbs sampler to infer signal transduction and (2) structural organization to uncover pathway landscape. Simulation studies reveal that GIP outperformed several existing approaches. We applied GIP to two tamoxifen-treated breast cancer patient data sets and identified robust signaling pathway networks associated with drug-resistance. Results were further validated using two drug-resistant cell line models to confirm the expression change in patent samples with or without drug resistance.

## Methods

Given predefined source proteins and target proteins, a flow network of given length is constructed between the sources and targets. Three potential functions, namely node potential, edge potential and flow potential, are defined for individual pathways to re-weight the flow network. Further, a probabilistic representation is formulated to convert pathway potential into a probability distribution. We apply a Gibbs sampling method to draw pathway samples according to the conditional distribution, which approximates the joint target distribution when the chain is sufficiently long. Finally, the algorithm highlights interconnected linear pathways with the largest potentials, where edge directions are inferred by aggregating pathway samples.

We define **θ**_1×*L*_ ={*θ*_1_, *θ*_2_, *θ*_*L*_} to denote a linear pathway of length *L*, where *θ*_*i*_, is a categorical variable that represents a specific protein at the *i*^th^ position of the pathway. *θ*_1_, is the source protein and *θ*_*L*_, is the target protein. Let Ω_*i*_ denote the domain of *θ*_*i*_, and we have 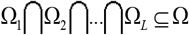, where Ω denotes all proteins in the PPI. We also have gene expression data **X**_*n*×*m*_, where *n* denotes number of genes and *m* denotes number of samples. We adopt the terminology of “potential function”, which originated from physics and was later widely used in image processing ^9^ to measure abnormality of pathways. We define three potential functions: node potential, edge potential and flow potential:

*Node potential* (V_1_ (*θ*_*i*_;**X**)): an aggregated z-score calculated from the discriminatory power of pathway members in differentiating two groups ^10^. Specifically, two-sample t-test is used to evaluate the differential expression of a member gene in the pathway. *P*-value is then calculated based on Student’s distribution of the t-statistic, and the aggregated z-score (node potential) is calculated using a mapping function defined by the normal inverse cumulative distribution function.
*Edge potential* (V_2_ (*θ*_*i*_,*θ*_*i*+1_;**X**)): an aggregated z-score calculated from the statistical significance of Pearson’s correlation between interacting proteins ^11^.
*Flow potential* (V_3_ (**θ**)): an z-score calculated based on the concordance between a pathway and its prior knowledge in order to incorporate the cellular location information.

We then combine the three potential functions as mentioned above linearly into a pathway energy function **θ** as follows:

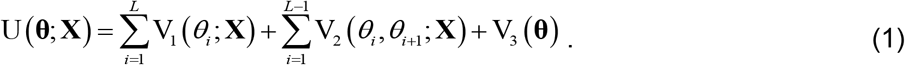

The first two potentials are driven by the data, which we refer to as “likelihood potentials”. The third flow potential comes from knowledge and is referred to as “prior potential”. We can obtain an optimal pathway by maximizing this function. However, there are many confounding factors including noise in gene expression data, tumor sample heterogeneity, false-positives and false-negatives in PPI, and the uncertainty of subcellular information induced by protein translocation. Simply finding the “best” pathway(s) cannot comprehensively capture signal transduction in the complex biological context of a cancer study. Therefore, we translate the optimization task into a distribution learning problem. We define an intermediate variable as S(**θ**) = −U(**θ**), and convert S(**θ**) into a probability using a Gibbs distribution ^12^ as defined in Eq. (2).

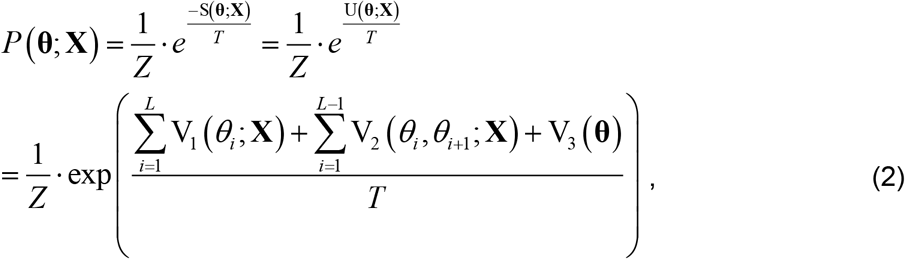

where 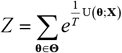 is a partition function and **Θ** is the set of all possible paths. *T* is the “temperature” that controls the shape of the distribution. To efficiently sample pathways from *P*(**θ**;**X**), we use a Gibbs strategy by iteratively drawing samples from a conditional distribution as follows:

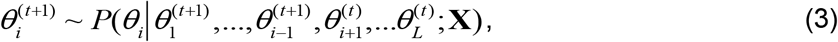

where *t* denotes the *t*^th^ sampling iteration. In each iteration, we accept probabilistically a new sample for *θ*_*i*_ based on the conditional distribution while other proteins *θ*_−*i*_ in the pathway are fixed. Samples drawn from this conditional distribution are theoretically good approximations of samples from the joint posteriors distribution.

To sample pathways of length *L*, we perform both forward and reverse searches from the source and target proteins, respectively. Consequently, we obtain a directed network structure of *L* layers (flow network) between the sources and targets. One protein may appear at different layers in the flow network. A flow network defines a regularized and directed structure between given source proteins and target proteins but uniform weights are assumed on each individual path. We re-weight the flow network by assigning potentials to individual nodes and edges. Starting from any arbitrary initial pathway, GIP iteratively replaces nodes in the *i*^th^ (1≤*i*≤*L*) layer of the current pathway by probabilistically sampling one candidate gene from the *i*^th^ layer.

After accumulating enough samples, GIP pools the samples together to estimate edge directions. We introduce a variable *e*_*i,j*_, where *e*_*i,j*_ = 1 denotes a directed edge from protien *ω*_*i*_∈Ω to protien *ω*_*i*_∈Ω. The underlying true probability of *e*_*i,j*_ is determined by the probability distribution of pathway **θ**, which is defined as:

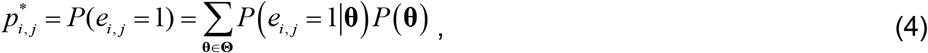

where *P*(*e*_*i,j*_ =1|**θ**)= 1 if *e*_*i,j*_ corresponds to a connected edge in pathway **θ** otherwise 0. Eq. (4) models the probability of each directed edge as a Bernoulli random variable with a success rate *p*_*i,j*_. A truncated version of *p*_*i,j*_ is also made available when users want to focus on top ranked pathways with large potentials (Supplementary Material Section A5). Now that we have calculated the probability of a directed edge (*e*_*i,j*_), we define the confidence of edge direction as:

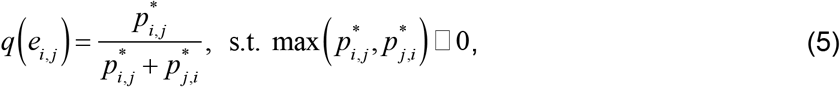

where *q*(*e*_*i,j*_)∊[0,1]. If *q*(*e*_*i,j*_) is close to 1, the signal flows from protein *ω*_*i*_ to *ω*_*j*_ with high confidence, while *q*(*e*_*i,j*_) = *q*(*e*_*i,j*_) = 0.5 indicates a lack of confidence in the direction of signal flow. Therefore, GIP not only assesses the importance of any individual edge, but also provides the confidence measure for edge direction.

## Results

### Performance evaluation with synthetic data

We first evaluated the performance of GIP for pathway identification on synthetic datasets generated by two basic pathway structures: type I structure and type II structure ^13^. Type I structure refers to wiring of alternative pathways between a single source protein and single target protein, while type II structure is established among multiple sources and targets. PPI data from the Human Protein Reference Database ^4^ and canonical pathways from KEGG ^14^ were downloaded to build simulation network and ground truth pathways. Table 1 summarizes the average precision ^15^ of our GIP method and three competing algorithms (random color coding ^16^, edge orientation ^8^, and integer linear programming (ILP) ^17^) on two simulation studies of type I and II structures under different noise settings. GIP has the best performance in identifying relevant pathway nodes and edges for both type I and type II structures.

**Table 1.**
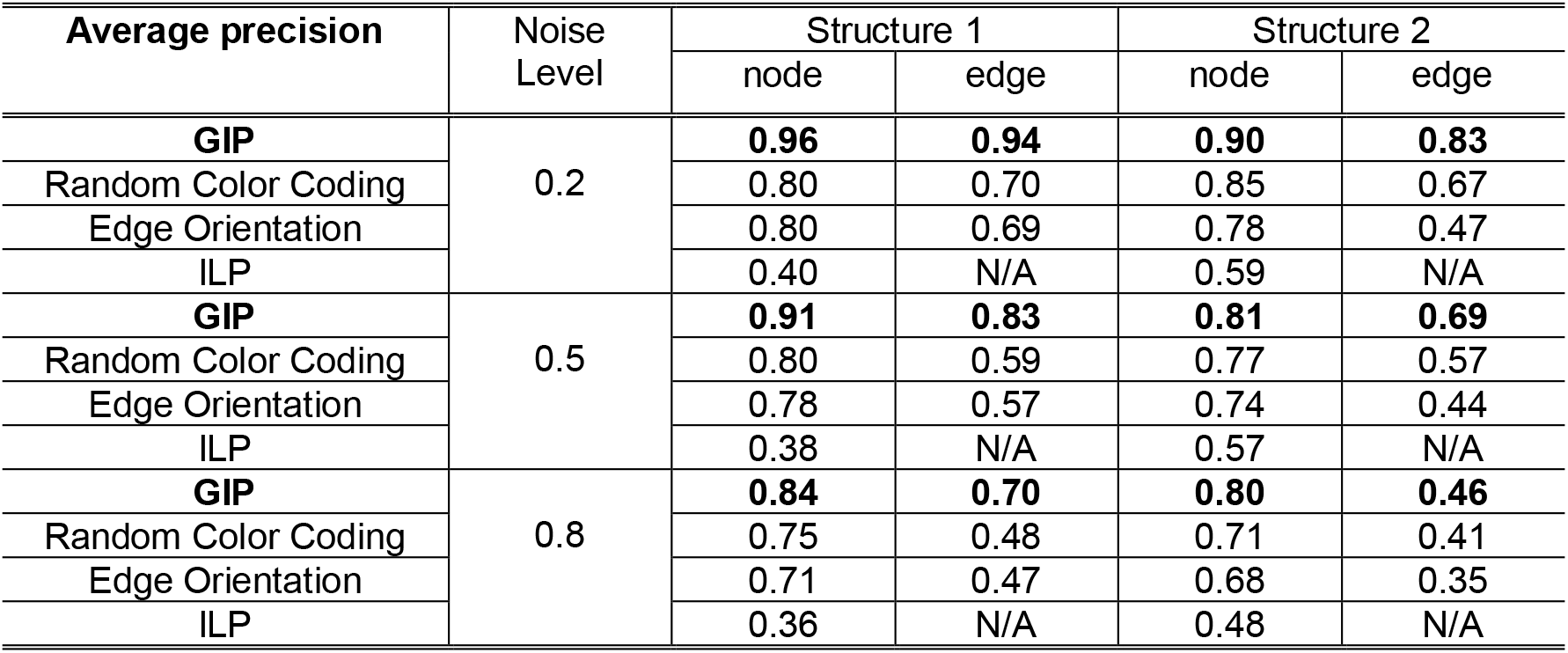
Average precision for pathway identification under different noise settings.

We also studied the impact of false positive connections in a PPI network on pathway identification. In Table 2, we summarized the average precision of all four methods for both type I and II pathway structures. It can be found that the performance of all four methods degrades as more false positive connections are introduced into the PPI network. However, GIP consistently maintains a comparable or better performance in all cases.

**Table 2.** Average precision for pathway identification under different proportion of false positive connections (PFC) in PPI network.

| Average precision | PFC | Structure 1 |  | Structure 2 |  |
| --- | --- | --- | --- | --- | --- |
|  |  | node | edge | node | Edge |
| **GIP** | 10% | **0.83** | **0.63** | **0.77** | **0.66** |
| Random Color Coding |  | 0.69 | 0.42 | 0.72 | 0.52 |
| Edge Orientation |  | 0.67 | 0.42 | 0.74 | 0.49 |
| ILP |  | 0.37 | N/A | 0.52 | N/A |
| **GIP** | 25% | **0.63** | **0.31** | **0.77** | **0.54** |
| Random Color Coding |  | 0.44 | 0.13 | 0.76 | 0.43 |
| Edge Orientation |  | 0.42 | 0.18 | 0.70 | 0.38 |
| ILP |  | 0.35 | N/A | 0.47 | N/A |
| **GIP** | 50% | **0.49** | **0.18** | **0.68** | **0.52** |
| Random Color Coding |  | 0.42 | 0.11 | 0.66 | 0.41 |
| Edge Orientation |  | 0.38 | 0.14 | 0.59 | 0.32 |
| ILP |  | 0.32 | N/A | 0.47 | N/A |

### Aberrant signal transduction identified from Tamoxifen treated patient data

We applied the GIP to a gene expression dataset ^18^ (termed Loi data here) to identify Tamoxifen resistance-related pathways in breast cancer. A 5-year cut-off on distant-metastasis-free-survival (DMFS) was used to divide samples into ‘early recurrence’ group (DMFS ≤ 5 years) and ‘late recurrence’ group (DMFS > 5 years), which finally yielded 88 ‘early recurrence’ samples and 92 ‘late recurrence’ samples in the Loi dataset ^19^. We used GIP to reconstruct signaling pathways related to estrogen receptor signaling, which connect source protein (ESR1) and functioning transcription factors identified using CRNET ^20^. An estrogen signaling pathway network between ESR1 and nuclear transcription factors was inferred by assembing the top 200 linear pathway samples. In total, there are 79 proteins in the pathway network, seven of which are transcription factors (JUN, FOS, STAT1, STAT3, STAT5A, ELK1, ETS1). We found complex wiring of alternative pathways going through frequently sampled cytoplasmic proteins of IRS1/2, JAK1, YWHAZ, CSNK2A1, MAPK1 and HSP90AA1. These proteins may play critical roles in exchanging messages among different functional modules; activate or suppress alternative pathways and/or initiate pathway crosstalk. We applied GIP to another Tamoxifen treated dataset ^21^ (termed Symmans data here) containing 47 ‘early recurrence’ samples and 56 ‘late recurrence’ samples using a 5-year survival cutoff. Results showed that for ER associated signaling pathway network, 72.5% (29/40) proteins identified from Symmans data overlapped with the proteins from Loi data.

### Validation using Tamoxifen-resistant breast cancer cell line model

We used breast cancer cell lines MCF7-STR, MCF7RR-STR, LCC1, LCC2 ^22^ growing *in vitro* to validate aberrant pathway modules identified from patient datasets, where MCF7RR-STR and LCC2 are Tamoxifen resistant cells, while MCF7-STR and LCC1 are antiestrogen sensitive. For ER signaling, a subset of genes and their expression levels were confirmed by both cell line studies including IRS1/2, FOS and JUN. Breast cancer relevant genes (BRCA1, BRCA2, CCNA2, E2F1, CDC25A, CDC25C, TOP2A, CDC2, CHUK) show dominant up-regulation in both the ‘early recurrence’ group of patient data and in the resistant cell line data, while transcription factors JUN, FOS, and STAT3 are down-regulated. We found that STMN1, PBK, CCNB1 and HSP90AA1 were overexpressed in early recurrence/resistant groups, while IRS1, IRS2, IGF1R and TSC2 were overexpressed in the late recurrence/non-resistant groups. The concordance between patient and cell line data supports the use of cell lines to study molecular signaling associated with the responsiveness to Tamoxifen and risk of breast cancer recurrence.

## Discussion

We have developed an integrative approach, GIP, to explore intracellular signal transduction and pathway landscape by integrating multi-omics data. GIP incorporate biological knowledge such as cellular location information, which makes the identified pathways more biologically interpretable. GIP also allow users to incorporate domain knowledge such as subcellular compartment by emphasizing signal transduction either in the nucleus, cytoplasm, or plasma membrane. The sampling framework has the felxibility to incorporate other genomic signals or other types of gene interactions to identify dysregulated signaling pathways. For example, many breast cancer genes carry somatic mutations ^23,24^. Such information can be integrated with mRNA expression to refine our pathway identification. Moreover, as transcription factors play a key role in signaling pathways and its binding to target gene is highly context-specific ^25,26^. Their functional interactions to target genes in a specific disease condition can facilitate disease-associated signaling pathway identification, too.

